# Sensitivity to sequential regularities in risky decision making

**DOI:** 10.1101/158253

**Authors:** Andrea Kóbor, Ádám Takács, Karolina Janacsek, Zsófia Kardos, Valéria Csépe, Dezso Nemeth

## Abstract

Probabilistic sequence learning involves a set of robust mechanisms that enable the extraction of statistical patterns embedded in the environment. It contributes to different perceptual and cognitive processes as well as to effective behavior adaptation, which is a crucial aspect of decision making. Although previous research attempted to model reinforcement learning and reward sensitivity in different risky decision-making paradigms, the basic mechanism of the sensitivity to statistical regularities has not been anchored to external tasks. Therefore, the present study aimed to investigate the statistical learning mechanism underlying individual differences in risky decision making. To reach this goal, we tested whether implicit probabilistic sequence learning and risky decision making share common variance. To have a more complex characterization of individual differences in risky decision making, hierarchical cluster analysis was conducted on performance data obtained in the Balloon Analogue Risk Task (BART) in a large sample of healthy young adults. Implicit probabilistic sequence learning was measured by the Alternating Serial Reaction Time (ASRT) task. According to the results, a four-cluster structure was identified involving average risk-taking, slowly responding, risk-taker, and risk-averse groups of participants, respectively. While the entire sample showed significant learning on the ASRT task, we found greater sensitivity to statistical regularities in the risk-taker and risk-averse groups than in participants with average risk-taking. These findings revealed common mechanisms in risky decision making and implicit probabilistic sequence learning and an adaptive aspect of higher risk taking on the BART. Our results could help to clarify the neurocognitive complexity of decision making and its individual differences.

## 1. Introduction

Decisions about skill-based actions are usually automatic in our daily routine, for instance, while we drive a car, do sports, or navigate in the operating system of our laptops. Making risky decisions, such as having one more drink before driving or driving a car beyond a speed limit, also involves automatic processes and gut feelings. We assume that decisions on both skill-based actions and risky situations necessitate the sensitivity to sequential or statistical regularities. An increased sensitivity to environmental regularities, which is also referred to as probabilistic sequence learning or statistical learning, has been shown to be crucial in healthy daily functioning, because it contributes to the acquisition of perceptual, motor, and cognitive skills, and to effective behavior adaptation (Batterink, Reber, Neville, & Paller, 2015; Chaudhuri & Fiete, 2016; Nemeth, Janacsek, & Fiser, 2013). Although probabilistic sequence learning and risky decision making could be connected, the majority of studies on risk-related behavior has focused on the effect of reward contingencies, task structure, different personality traits and/or clinical symptoms (e.g., Bornovalova et al., 2009; Brand, Labudda, & Markowitsch, 2006; Fein & Chang, 2008; Schiebener, Wegmann, Pawlikowski, & Brand, 2012), and the direct investigation of the association between this learning mechanism and risk-related behavior has been missing from the field. Learning per se has been linked to the adaptive nature of risk-related behavior (Bechara, Damasio, Tranel, & Damasio, 2005; Euser et al., 2013), but the involvement of the ability to acquire statistical contingencies has been unclear. Here we aim to fill in this gap by analyzing task-solving strategies during a sequential risk-taking task with probabilistic underlying structure in a large population-based sample and testing its relations to probabilistic sequence learning measured by an independent perceptual-motor four-choice reaction time task that involves implicit learning of statistical regularities.

The Balloon Analogue Risk Task (BART) has been considered as a valid measure of naturalistic risk-taking behavior by modeling the day-to-day sequential processing of risk (Helfinstein et al., 2014; Lejuez et al., 2002; Schonberg et al., 2012; Schonberg, Fox, & Poldrack, 2011). In this task, participants are asked to inflate an empty virtual balloon. Each balloon pump is associated with either a reward or a balloon burst. The probability of a balloon burst increases with each successive pump, but the regularity that determines balloon bursts following a pump is unknown to participants. In previous experiments, probabilities of balloon bursts have usually been chosen from a uniform distribution, and participants have had to infer these probabilities by trial and error learning (Schonberg et al., 2011). As the appearance of gains (balloon increase) and losses (balloon burst) follows a probabilistic structure, expectations about stimulus-response contingencies might not be established on the basis of purely explicit task-solving strategies.

Previous results also suggest that the sequential processing of risk in the BART evokes expectations about outcome contingencies (Kardos et al., 2016; Kiat, Straley, & Cheadle, 2016), and the process of expectation formation is a crucial element of probabilistic sequence learning, as well. According to this argument, the BART might share common variance with other external measures of probabilistic sequence learning. This assumption also follows from another line of research suggesting interactions between different learning/memory systems (for a review, see Robertson, 2012) and learning transfer between learning/memory tasks (Mosha & Robertson, 2016). However, it has also been shown that multiple task-related sequential regularities could be acquired in parallel (Goschke & Bolte, 2012), based on which the lack of relation between learning two different dimensions of sequences (i.e., the sequence of spatial positions in the perceptual-motor task and the sequence of response-outcome probabilities in the BART) is also possible.

As the BART is a widely used tool in the clinical and experimental literature (Lauriola, Panno, Levin, & Lejuez, 2014), previous work has already attempted to go beyond the common behavioral variables to describe performance and thus capture more processes of task solving. Schmitz, Manske, Preckel, and Wilhelm (2016) differentiated variables related to risk taking, task performance, impulsive decision making, and reinforcement sequence modulation. Using a shortened version of the BART, on the basis of two empirical studies on adolescents and young adults (among whom a significant proportion showed deviant behavior), the authors suggested that the number of balloon bursts was the most consistent correlate of risk taking with a high predictive validity, and the use of RT-based scores indicating impulsive decision making was limited.

With the same purpose, to more clearly characterize the underlying psychological/cognitive processes determining BART performance, another line of research has focused on developing formal models of task. Wallsten, Pleskac, and Lejuez (2005) compared alternative models of the BART and found that a four-parameter model provided the best fit to data. This model indicated that the decision makers assumed stationary burst probabilities over pumps, they learned - updated their opinion about burst probabilities - in a Bayesian fashion over balloon trials, their initial risk preferences were evaluated prior to responding, and their response consistency remained constant over trials. The study of Pleskac, Wallsten, Wang, and Lejuez (2008) added to these findings by providing evidence that decision makers adapted their mental representation and learning processes according to the actual stochastic structure of the decision task when a modified version of the BART was applied. It was also suggested that the ill-defined, nonstationary characteristic of the original task, which was related to learning processes, hindered its predictive validity to identify real-world risk taking behavior. Meanwhile, on data derived from a BART version with fixed bursting probability over trials, van Ravenzwaaij, Dutilh, and Wagenmakers (2011) found that a simplified, two-parameter (risk taking and response consistency) version of the model introduced by Wallsten et al. (2005) showed adequate parameter recovery instead of the four-parameter model.

The advantage of cognitive modeling over analyzing the standard behavioral variables of the BART has been shown, for instance, in the study of Rolison, Hanoch, and Wood (2012), where no difference was found between younger and older adults in risky behavior according to the standard BART score, but modeling results revealed that older adults were initially more risk averse and then adjusted their behavior according to experience. Similarly, differences in those psychological processes that model parameters represent were found in the study of Wichary, Pachur, Kościelniak, Rydzewska, and Sedek (2017) between young and older adults experiencing initial good and bad luck in the BART (see also Koscielniak, Rydzewska, & Sedek, 2016). In addition, Wichary, Pachur, and Li (2015) revealed striking gender differences in model parameters between individuals with excessive risk taking (prisoners) and control participants.

Although using other analytic approaches, learning in the BART has also been quantified by tracking how participants increase the number of balloon pumps after they have gained some experience with the task during earlier balloon trials (Campbell, Samartgis, & Crowe, 2013; Euser et al., 2013; Euser, van Meel, Snelleman, & Franken, 2011; Fecteau et al., 2007; Koscielniak et al., 2016; Lim, Yuen, & Tong, 2015; Vigil-Colet, 2007), and how they change their behavior on a particular trial according to the outcome on the preceding trial (Courtney et al., 2012; Kohno et al., 2015). In addition, an increasing number of studies has focused on how response time of participants could indicate the change of decision-making processes throughout the BART (e.g., Euser et al., 2011; Hassall, Holland, & Krigolson, 2013; Pleskac & Wershbale, 2014; Schonberg et al., 2012). These studies quantified different decision making mechanisms within the BART; however, these mechanisms have not been assigned to external measures of implicit acquisition of statistical regularities that could characterize sequential decision making.

While the above-mentioned studies have been promising in understanding the cognitive processes underlying the BART, it is not clear whether the *combination* of basic behavioral indices might describe distinctive task-solving profiles, which could further promote the investigation of individual differences in risky decision making. Specifically, more accurate task-solving profiles could contribute to revealing the exact relationship between probabilistic sequence learning and BART performance, which otherwise would have remained hidden. For a better characterization of individual differences in risky decision making, classification of participants on the basis of their behavioral performance using different clustering methods seems to be an advantageous approach (Bergman, Magnusson, & El-Khouri, 2003; Kóbor, Takács, Urbán, & Csápe, 2012). Therefore, in this study, we performed hierarchical cluster analysis to capture individual differences in BART performance. In two steps, we tested whether the BART and an implicit probabilistic sequence learning task share common variance as both tasks involve probabilistic underlying structure. First, we checked the potential associations between the component measures of the BART and probabilistic sequence learning. Second, the sequence learning performance of the clusters of participants with different task-solving strategies were compared.

## 2. Material and Methods

### 2.1 Participants

The sample consisted of 180 healthy young adults. Mainly the undergraduate students of Eötvös Loránd University participated in this study. Descriptive characteristics of the sample are presented in Table 1 (see the column labeled as “Total sample”). All participants had normal or corrected-to-normal vision and none of them reported a history of any neurological and/or psychiatric condition. All participants provided written informed consent before enrolment and received course credits for taking part in the experiment. The study was approved by the United Ethical Review Committee for Research in Psychology (EPKEB) in Hungary (approval number: 30/2012). The study was conducted in accordance with the Declaration of Helsinki.

**Table 1.**
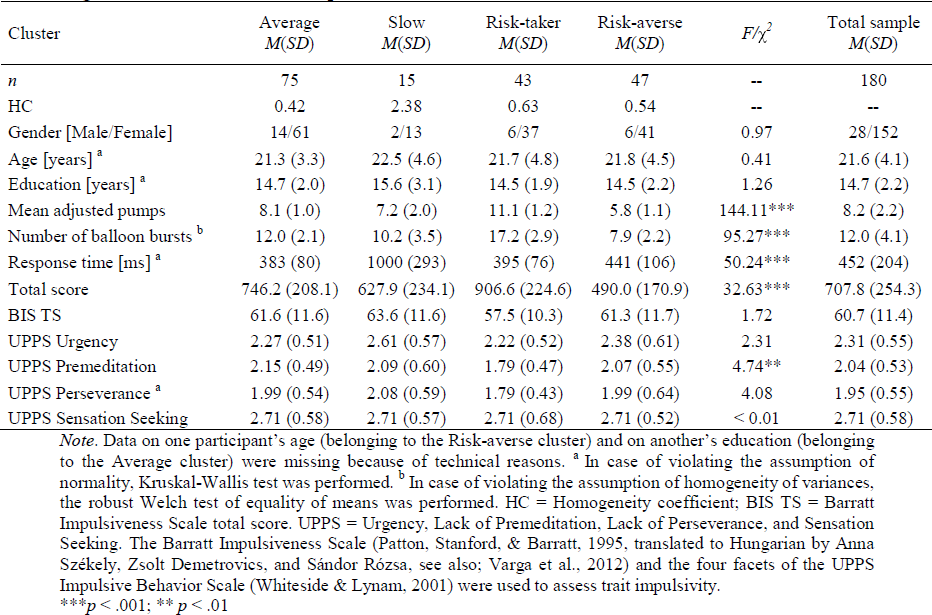
Descriptive data of demographic variables, BART performance, and rating scales in the four strategic clusters and the total sample.

| Cluster | Average<br>*M(SD)* | Slow<br>*M(SD)* | Risk-taker<br>*M(SD)* | Risk-averse<br>*M(SD)* | $F/\chi^2$ | Total sample<br>*M(SD)* |
| --- | --- | --- | --- | --- | --- | --- |
| *n* | 75 | 15 | 43 | 47 | -- | 180 |
| HC | 0.42 | 2.38 | 0.63 | 0.54 | -- | -- |
| Gender [Male/Female] | 14/61 | 2/13 | 6/37 | 6/41 | 0.97 | 28/152 |
| Age [years] <sup>a</sup> | 21.3 (3.3) | 22.5 (4.6) | 21.7 (4.8) | 21.8 (4.5) | 0.41 | 21.6 (4.1) |
| Education [years] <sup>a</sup> | 14.7 (2.0) | 15.6 (3.1) | 14.5 (1.9) | 14.5 (2.2) | 1.26 | 14.7 (2.2) |
| Mean adjusted pumps | 8.1 (1.0) | 7.2 (2.0) | 11.1 (1.2) | 5.8 (1.1) | 144.11*** | 8.2 (2.2) |
| Number of balloon bursts <sup>b</sup> | 12.0 (2.1) | 10.2 (3.5) | 17.2 (2.9) | 7.9 (2.2) | 95.27*** | 12.0 (4.1) |
| Response time [ms] <sup>a</sup> | 383 (80) | 1000 (293) | 395 (76) | 441 (106) | 50.24*** | 452 (204) |
| Total score | 746.2 (208.1) | 627.9 (234.1) | 906.6 (224.6) | 490.0 (170.9) | 32.63*** | 707.8 (254.3) |
| BIS TS | 61.6 (11.6) | 63.6 (11.6) | 57.5 (10.3) | 61.3 (11.7) | 1.72 | 60.7 (11.4) |
| UPPS Urgency | 2.27 (0.51) | 2.61 (0.57) | 2.22 (0.52) | 2.38 (0.61) | 2.31 | 2.31 (0.55) |
| UPPS Premeditation | 2.15 (0.49) | 2.09 (0.60) | 1.79 (0.47) | 2.07 (0.55) | 4.74** | 2.04 (0.53) |
| UPPS Perseverance <sup>a</sup> | 1.99 (0.54) | 2.08 (0.59) | 1.79 (0.43) | 1.99 (0.64) | 4.08 | 1.95 (0.55) |
| UPPS Sensation Seeking | 2.71 (0.58) | 2.71 (0.57) | 2.71 (0.68) | 2.71 (0.52) | < 0.01 | 2.71 (0.58) |

Note. Data on one participant's age (belonging to the Risk-averse cluster) and on another's education (belonging to the Average cluster) were missing because of technical reasons. <sup>a</sup> In case of violating the assumption of normality, Kruskal-Wallis test was performed. <sup>b</sup> In case of violating the assumption of homogeneity of variances, the robust Welch test of equality of means was performed. HC = Homogeneity coefficient; BIS TS = Barratt Impulsiveness Scale total score. UPPS = Urgency, Lack of Premeditation, Lack of Perseverance, and Sensation Seeking. The Barratt Impulsiveness Scale (Patton, Stanford, & Barratt, 1995, translated to Hungarian by Anna Székely, Zsolt Demetrovics, and Sándor Rózsa, see also; Varga et al., 2012) and the four facets of the UPPS Impulsive Behavior Scale (Whiteside & Lynam, 2001) were used to assess trait impulsivity.
\*\*\* $p < .001$ ; \*\* $p < .01$

### 2.2 Tasks

#### 2.2.1 BART

The general structure and appearance of the BART was the same as described in previous studies (Fein & Chang, 2008; Kóbor et al., 2015; Takács et al., 2015). Participants were instructed to collect as many points as possible by inflating an empty virtual balloon on a screen. Accumulated score for a given balloon, which simultaneously increased with the size of the balloon after each successful pump, were displayed in the middle of the balloon. Two response keys on a keyboard were designated either to pump (Z) the balloon or to finish the actual trial and collect (C) the accumulated score. Instead of collecting the score from the actual balloon, there were two possible outcomes as a results of a further pump: An increase in the size of the balloon together with an increase in the score inside (positive feedback) or a balloon burst (negative feedback) could have happened. The balloon burst ended the actual trial, and the accumulated score on that balloon was lost.

Importantly, each successful pump increased the probability of a balloon burst and the accumulated score being lost. The regularity determining balloon bursts was unknown to participants and followed three principles: (1) balloon bursts for the first and second pumps were disabled; (2) the maximum number of successful pumps for each balloon was 19; (3) the probability of a balloon burst was 1/18 for the third pump, 1/17 for the fourth pump, and so on for each further pump until the 20^th^, where the probability of a balloon burst was 1/1. Compared to the typical variant of the task (Lejuez et al., 2002), we modified the increase of payoffs to motivate participants to take higher risk and gain more reward (cf. Fein & Chang, 2008). Namely, we assumed that because of the higher appealing characteristic of reward, participants would be more prone to test the structure of the task. Therefore, reward score increased by one point at each successful pump: Participants could gain one point for the first pump, two for the second (i.e., the accumulated score for a given balloon was three), three for the third (i.e., the accumulated score was six), and so on. Our previous studies (Kardos et al., 2016; Kóbor et al., 2015; Takács et al., 2015) indicate that behavioral results have been similar in this variant of the task to that of the typical variant (Lejuez et al., 2002).

In the middle of the balloon, participants always saw the total accumulated score for a given balloon. The labels “Total score” depicting the points in the permanent bank, “Last balloon” depicting the points collected from the previous balloon, and response key options constantly appeared on the screen during the experiment. After collecting the accumulated reward, a separate screen indicated the gained score. This screen or the other presenting balloon burst was followed by the presentation of a new empty (small-sized) balloon indicating the beginning of the next trial.

In this version of the BART, participants had to inflate 30 balloons. In order to maximize reward, the optimal or advantageous number of pumps was 13, but participants had to infer this information by trial-and-error learning. Therefore, approaching this particular value by increasing the number of pumps in time could be regarded as the evolvement of sensitivity to the underlying statistical regularities.

#### 2.2.2 ASRT task

The Alternating Serial Reaction Time (ASRT) task was used to measure implicit probabilistic sequence learning (Nemeth et al., 2010). In this task, the target stimulus was a picture of a dog’s head, which appeared in one of four horizontally arranged and equally spaced empty circles on the screen in each trial (Nemeth, Janacsek, Polner, & Kovacs, 2013). Participants were instructed to press a key (Z, C, B, or M on a QWERTY keyboard) corresponding to target location as quickly and accurately as they can. The target stimulus remained on the screen until the participants’ correct response, and the next target was presented on the screen after 120 ms delay. Unbeknownst to the participants, the presentation of stimuli followed an eight-element sequence, within which pattern (P) and random (r) elements alternated with each other (e.g., 2 - *r* - 1 - *r* - 3 - *r* - 4 - *r*; where numbers denote the four locations on the screen from left to right, and r’s denote randomly chosen locations out of the four possible ones). In each block, this eight-element trial sequence was repeated 10 times after five warmup trials consisting only of random stimuli (altogether 85 trials in each block).

As a results of the trial sequence, some patterns of three successive elements (henceforth referred to as triplets) occur more frequently than others in the ASRT task. In the example above, 2X1, 1X3, 3X4, and 4X2 (X indicates the middle element of the triplet) occurred often since their third elements could have either been a pattern or a random element. However, 1X2 and 4X3 occurred less frequently since their third element could have only been random. The former triplet types were labeled as “high-frequency” triplets while the latter types were labeled as “low-frequency” triplets (Nemeth, Janacsek, & Fiser, 2013). The third element of a high-frequency triplet was more predictable from the first element of the triplet than in the case of low-frequency triplets. Accordingly, each target stimulus was categorized as either the third element of a high - or a low-frequency triplet, and the accuracy and reaction time (RT) of the response to this item were compared between the two triplet types.

While high frequency triplets could be expected with 62.5% of probability, low frequency triplets had a 37.5% probability to occur. Following the standard analysis protocol of previous studies (J. H. Howard, Jr. & Howard, 1997; Nemeth, Janacsek, & Fiser, 2013), two types of low-frequency triplets were eliminated from the analysis: repetitions (e.g., 111, 444) and trills (e.g., 121, 242). Repetitions and trills were low frequency for all participants, and participants often show pre-existing response tendencies to them (D. V. Howard et al., 2004). By eliminating these triplets, we could ensure that any high-versus low-frequency differences were due to learning and not to pre-existing tendencies. Probabilistic sequence learning is reflected in the increasingly faster and more accurate responses to high-frequency triplets compared to that to low-frequency ones over the course of the task (S. Song, J. H. Howard, Jr., & D. V. Howard, 2007b). In addition, it has been shown that accuracy decreases on low-frequency (less predictable) triplets as a results of probabilistic sequence learning (D. V. Howard et al., 2004). Consequently, the obtained learning measure could also be considered as an index of probabilistic sequence learning. It is important to note that the task remained implicit for the participants, and according to previous studies, even after an extended practice, participants were not able to discover the hidden sequence (D. V. Howard et al., 2004).

### 2.3 Procedure

The ASRT task consisted of 45 blocks with 85 trials in each. Participants were allowed to take a short break between blocks. In a separate experimental session 24 hours after completing the ASRT task, we administered the BART, other neuropsychological tests, and questionnaires measuring the different aspects of cognition, personality, and social behavior. Here we only report results of the BART and the ASRT task.

### 2.4 Statistical Analyses

#### 2.4.1 BART variables

We followed a theory-driven approach (Appelt, Milch, Handgraaf, & Weber, 2011) as well as considered the most frequently published behavioral indices of the BART when deciding about the individual component measures characterizing BART performance. The choice of these variables is also critical in regard to the obtainable cluster solution (Morris, Blashfield, & Satz, 1981). Three variables were determined. First, the *mean adjusted number of pumps* across balloons (MAP; mean number of pumps on balloons that did not burst) is conventionally used to measure risk-taking behavior (Lejuez et al., 2002), and it could also indicate how participants learn from positive feedback. In other words, higher MAP could mirror a more optimal task-solving strategy. Second, the *number of balloon bursts* (i.e., pop number) not only indicates the level of risk taking (Schmitz et al., 2016) but also the effect of negative feedback throughout the task. Besides general risk-taking behavior and insensitivity to losses, higher pop number could mirror higher propensity to test the structure of the task and a step towards optimal task-solving strategy. In addition, it seems that participants pay more attention to losses than to gains (Rolison et al., 2012). According to Schmitz et al. (2016), the number of balloon bursts has been a less ambivalent indicator of risk taking than the MAP. Therefore, in this study, we consider variability in the MAP as an indicator of variability in optimal task solving, while variability in the number of balloon bursts is assumed to be related to variability in risk taking and optimal task solving, as well. The third variable we used was the *median response time (RT) of pumps* across balloons calculated for each participant. Response time was measured from the presentation of the (empty or increased) balloon until the initiation of the next pump. Balloon pumps with RT equal to or below 100 ms and equal to or higher than 3000 ms were excluded from calculating the median of all RTs in order to eliminate attentional lapses and premature responses (cf. Hassall et al., 2013; Kardos et al., 2016; Matzke & Wagenmakers, 2009). According to previous studies (Hassall et al., 2013; Pleskac & Wershbale, 2014; Wallsten et al., 2005), RT of balloon pumps could be indicative of how participants explore the reward structure of the task and make risk assessment before each decision. Basically, generally slower RTs could be related to more controlled, deliberate pumps and the exploration of reward contingencies throughout the task (Haffke & Hübner, 2015; Pleskac et al., 2008).

Although the overall quality (effectiveness) of decision making as well as the adaptation to task requirements (in line with task instructions) could be captured by the *total score* (Koscielniak et al., 2016; Schmitz et al., 2016), we did not use this variable for clustering. A certain value on this variable accumulates the outcome of many different processes and strategies underlying decision making, and previously, it has been related to scholastic achievement and working memory but not to risk-taking variables (Schmitz et al., 2016). As we intended to characterize performance by combining different pieces of information conveyed by each component measure of the BART, we regarded the total score only as a measure to externally validate our cluster solution. An appropriate cluster solution should show differences on a related overall performance variable (i.e., the total score) that has not been used for clustering (Morris et al., 1981).

#### 2.4.2 ASRT task performance

Statistical analyses of the ASRT performance followed the protocol established in previous studies (J. H. Howard, Jr. & Howard, 1997; Romano, Howard, & Howard, 2010). Five-block-long segments of data were collapsed into larger epochs; thus, we altogether analyzed 9 epochs of the ASRT task. Epochs are labeled consecutively in this paper (1, 2, etc.). For each participant and epoch, we calculated mean accuracy (percentage of correct responses) and median RT (only for correct responses), separately for high-and low-frequency triplets. Then we calculated a learning score as the difference between triplet types in RT (RT for low-frequency triplets minus RT for high-frequency triplets) and accuracy (accuracy for high-frequency triplets minus accuracy for low-frequency triplets). Larger score in both measures indicates larger probabilistic sequence learning.

#### 2.4.3 The association between BART variables and ASRT task performance

In order to check whether any variability is shared between the two, theoretically related functions, statistical analysis was performed in two steps. First, we calculated Pearson’s linear correlations between the component measures of the BART and ASRT learning scores. Second, as a more fine-grained characterization of performance, for identifying subgroups of distinctive task-solving strategies, we performed an agglomerative hierarchical cluster analysis with the clustering variables of MAP, number of pops, and median RT of balloon pumps. We used squared Euclidean distance as the similarity measure and Ward’s method as the type of cluster fusion (Morey, Blashfield, & Skinner, 1983). Before performing the cluster analysis, the three clustering variables were standardized (they were transformed into *z*scores). After the hierarchical cluster analysis, to further improve the obtained cluster solution and create more homogeneous subgroup, we performed *K*-means cluster analysis. This iterative method moves (relocates) some cases from one cluster to another if this reduces the total error sum of squares of the original cluster solution (Bergman et al., 2003; Takács, Kóbor, Tárnok, & Vargha, 2014). To evaluate probabilistic sequence learning and compare probabilistic sequence learning performance of subgroups obtained from the final cluster solution, mixed design analyses of variance (ANOVAs) were conducted. Greenhouse-Geisser epsilon (ε) correction was used when necessary. Original *df* values and corrected*p* values (if applicable) are reported together with partial eta-squared (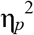) as the measure of effect size. We used LSD (Least Significant Difference) tests for pair-wise comparisons.

## 3. Results

### 3.1 Change in behavioral performance during the BART

To check whether participants have tried to optimize their performance during task solving, we analyzed the change in behavior over time. We calculated the MAP and the number of balloon bursts for the first, second, and third 10 balloons, respectively, for the whole sample. First, a one-way repeated measures ANOVA with BIN (1-10, 11-20, 21-30 balloons) as a within-subjects factor was performed on the MAP. The main effect of BIN was significant, *F* (2, 358) = 38.37, ε = .883,*p* < .001, 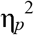 = .177. Pair-wise comparisons showed that the MAP gradually increased in the whole sample as the task progressed (*M*_first bin_ = 7.4, *M*_second bin_ = 8.5, *M*_hird bin_ = 8.9, all comparisons are significant, *ps* ≤ .008).

Second, the same ANOVA was performed on the number of balloon bursts. The main effect of BIN was significant, *F* (2, 358) = 19.34, *p* < .001, 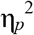 = .098. Pair-wise comparisons showed that the number of balloon bursts increased from the first to the second and third bins (*M*_first bin_ = 3.5, *M*_second bin_ = 4.2, *M*_third bin_ = 4.4; first vs. second: *p* < .001; first vs. third: *p* < .001), but there was no difference between the second and third bins (*p* = .172). These finding suggest that participants were sensitive to statistical regularities underlying the BART as they tried to test the structure of the task, at least during the first 20 balloons.

### 3.2 BART clusters

Cluster analysis was performed using the overall MAP, number of balloon bursts, and response time variables, calculated across 30 balloons. The final cluster solution included four clusters explaining 67.58% of the variance (considering error sum of squares). The Silhouette coefficient of the cluster solution was .681. This coefficient indicates the quality of cluster cohesion and separation, it ranges between -1 and 1, and values greater than .5 indicates reasonable partitioning of the data. The average of the Homogeneity coefficient (HC) was 0.667. HC is the average of the pairwise distances within a cluster; larger values indicate more heterogeneous clusters, and an average HC less than 1 indicates good cluster structure. (For more details on evaluating cluster solutions, see Vargha, Bergman, and Takács (2016)).

Detailed demographic and behavioral properties of the four clusters and the entire sample are presented in Table 1 and Figure 1. We labeled and interpreted the clusters on the basis of their descriptive characteristics shown on BART behavioral measures and following the notion that higher MAP and higher number of balloon bursts could indicate more optimal, while higher RT could indicate more deliberate task solving. Accordingly, the first cluster involved participants with moderate or average risk-taking (41.7%), the second cluster captured slowly responding participants (8.3%), the third cluster consisted of risk-taker participants (23.9%), and the fourth cluster consisted of risk-averse ones (26.1%). Participants’ mean values on BART outcome measures in the Average cluster were close to that of the total sample, except the RT, which was slightly faster. Slowly responding participants experienced relatively low number of balloon bursts and produced relatively low MAP. Number of balloon bursts and the MAP were even lower in the Risk-averse cluster, which was otherwise described by average RTs. The mean of MAP in the Risk-taker cluster was closer to the optimal level than in other clusters, and participant in this subgroup also experienced a high number of balloon bursts. According to pair-wise comparisons, each cluster differed from all the others on the total score: The Risk-taker cluster achieved the highest total score, the Average cluster was the second, the Slow cluster was the third, and the Risk-averse cluster achieved the lowest total score (all *ps* ≤ .043).

**Figure 1.**
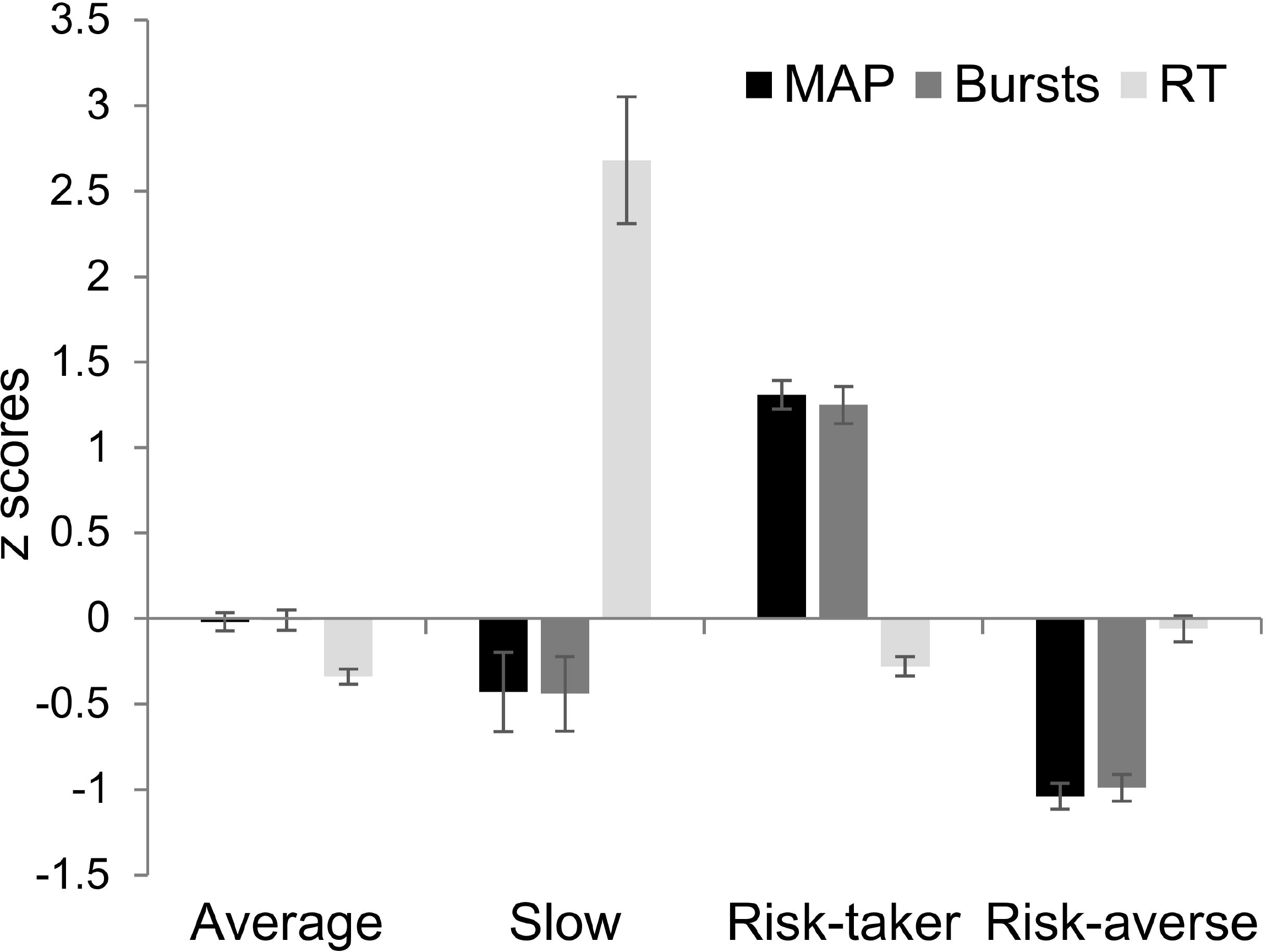
Behavioral profiles of the four clusters on the three BART variables used for clustering. Error bars denote standard error of mean. MAP = mean adjusted number of pumps, Bursts = Number of balloon bursts, RT = Response time.

### 3.3 Associations between the BART and the ASRT task

One participant was removed from the following analyses because of high error ratio on the ASRT task (a mean of 74% for the entire task): This case was an extreme outlier, being well below the lower whisker of the sample’s accuracy data represented as a boxplot (sample’s accuracy: *M* = 95.40%, *SD* = 2.95%). Therefore, *n* = 179 in the remainder of the paper.

#### 3.3.1 Correlation analysis

Regarding the whole sample, there was *no significant* correlation between either the accuracy learning score of the ASRT task and BART measures (MAP: *r* = .002, *p* = .983; pop: *r* = -.013, *p* = .864; RT: *r* = -.019, *p* = .798; total score: *r* = -.011, *p* = .881; *df* = 177 in all analyses), or the RT learning score of the ASRT task and BART measures (MAP: *r* = -.069, *p* = .361; pop: *r* = -.090, *p* = .232; RT: *r* = -.065, *p* = .386; total score: *r* = -.052, *p* = .491; *df=*177 in all analyses). We also plotted each learning index against each component measure of the BART, and no indication was found for linear or quadratic relations (for the sake of brevity, these figures are not included).

#### 3.3.2 Between-cluster differences on the ASRT task

Learning on the ASRT task among the BART strategic clusters was tested with a three-way mixed ANOVA on *accuracy* with TRIPLET (high-vs. low-frequency) and EPOCH (1-9) as within-subjects factors and CLUSTER (Average, Slow, Risk-taker, Risk-averse) as a between-subjects factor. Accuracy data as a function of epoch and trial type for each cluster are shown in Figure 2. We first present the task-related (within-subjects) effects. The significant main effect of TRIPLET, *F* (1, 175) = 344.55,*p* < .001, 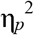 = .663, revealed probabilistic sequence learning in the entire sample since participants were less accurate on low-frequency triplets than on high-frequency triplets. The significant main effect of EPOCH, *F* (8, 1400) = 30.55, ε = .537, *p* < .001, 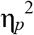 = .149, also indicated learning, as it changed mainly due to the decreasing accuracy on low-frequency triplets throughout the task (cf. D. V. Howard et al., 2004). Specifically, participants became less accurate on low-frequency triplets than on high-frequency ones as the task progressed, reflected by the significant interaction of TRIPLET*EPOCH, *F* (8, 1400) = 14.76, ε = .904, *p* < .001, 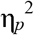 = .078.

**Figure 2.**
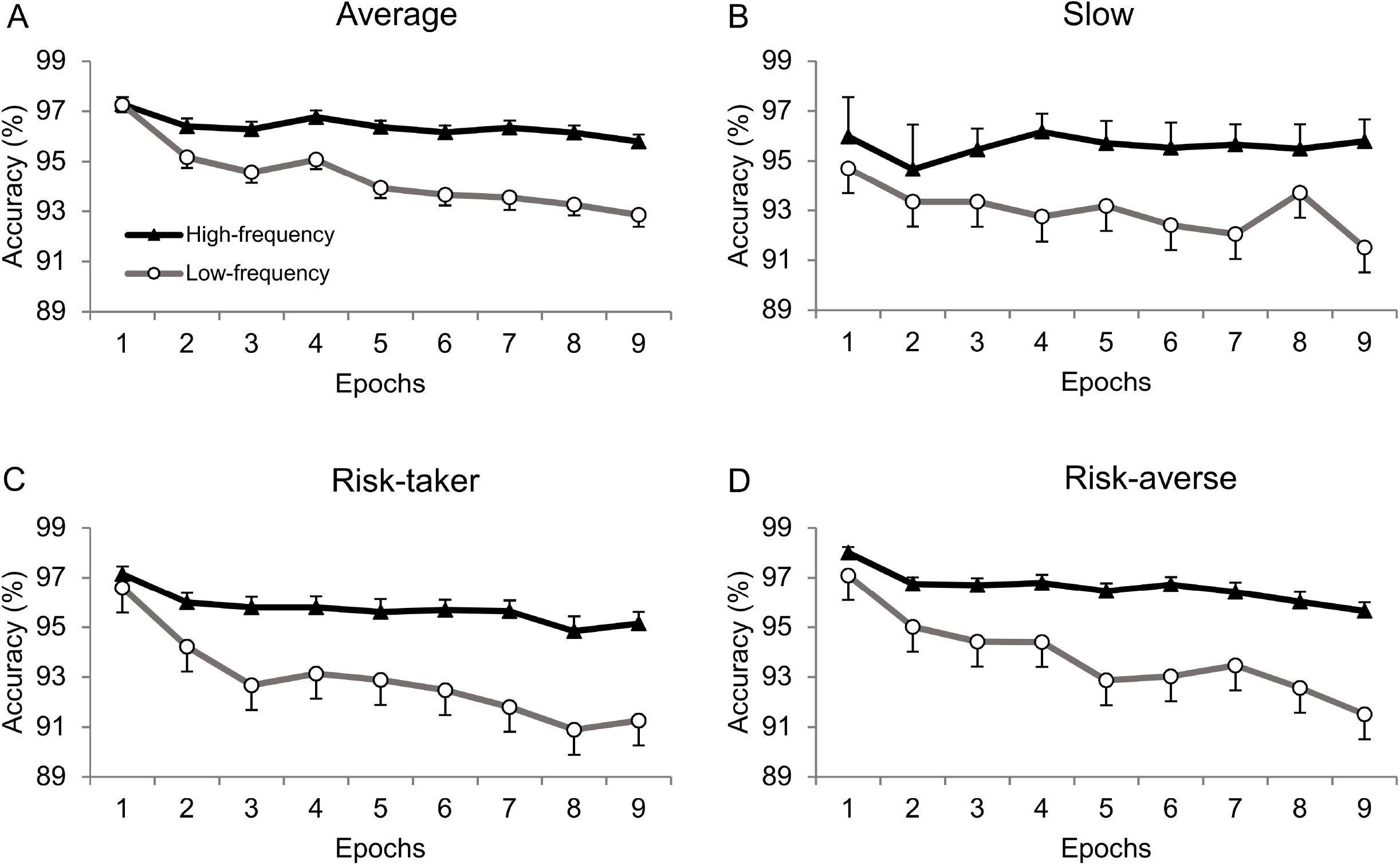
Temporal dynamics of probabilistic sequence learning. Data are presented on the accuracy measure across epochs (1-9) as a function of trial type (high-vs. low-frequency triplets), separately for each cluster. (A) Average, (B) Slow, (C) Risk-taker, (D) Risk-averse. Error bars denote standard error of mean.

In regard to the between-subjects effects on ASRT accuracy measures, the main effect of CLUSTER did not reach significance, *F* (3, 175) = 1.91, *p* = .131, 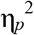 = .032, showing that overall accuracy did not reliable differ between the BART clusters. However, the TRIPLET*CLUSTER interaction was significant, *F* (3, 175) = 3.48, *p* = .017, 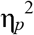 = .056, indicating differences between the BART clusters in probabilistic sequence learning. Learning score was greater in the Risk-taker and Risk-averse clusters than in the Average cluster (*ps* < .010, see Figure 3). The EPOCH*CLUSTER interaction only tended to be significant, *F* (24, 1400) = 1.57, ε = .357, *p* = .089, 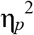 = .026. Nevertheless, the TRIPLET*EPOCH*CLUSTER interaction was not significant, *F* (24, 1400) = 0.784, ε = .904,*p* = .747, 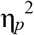 = .013, suggesting that the time course of probabilistic sequence learning was similar across the BART clusters.

**Figure 3.**
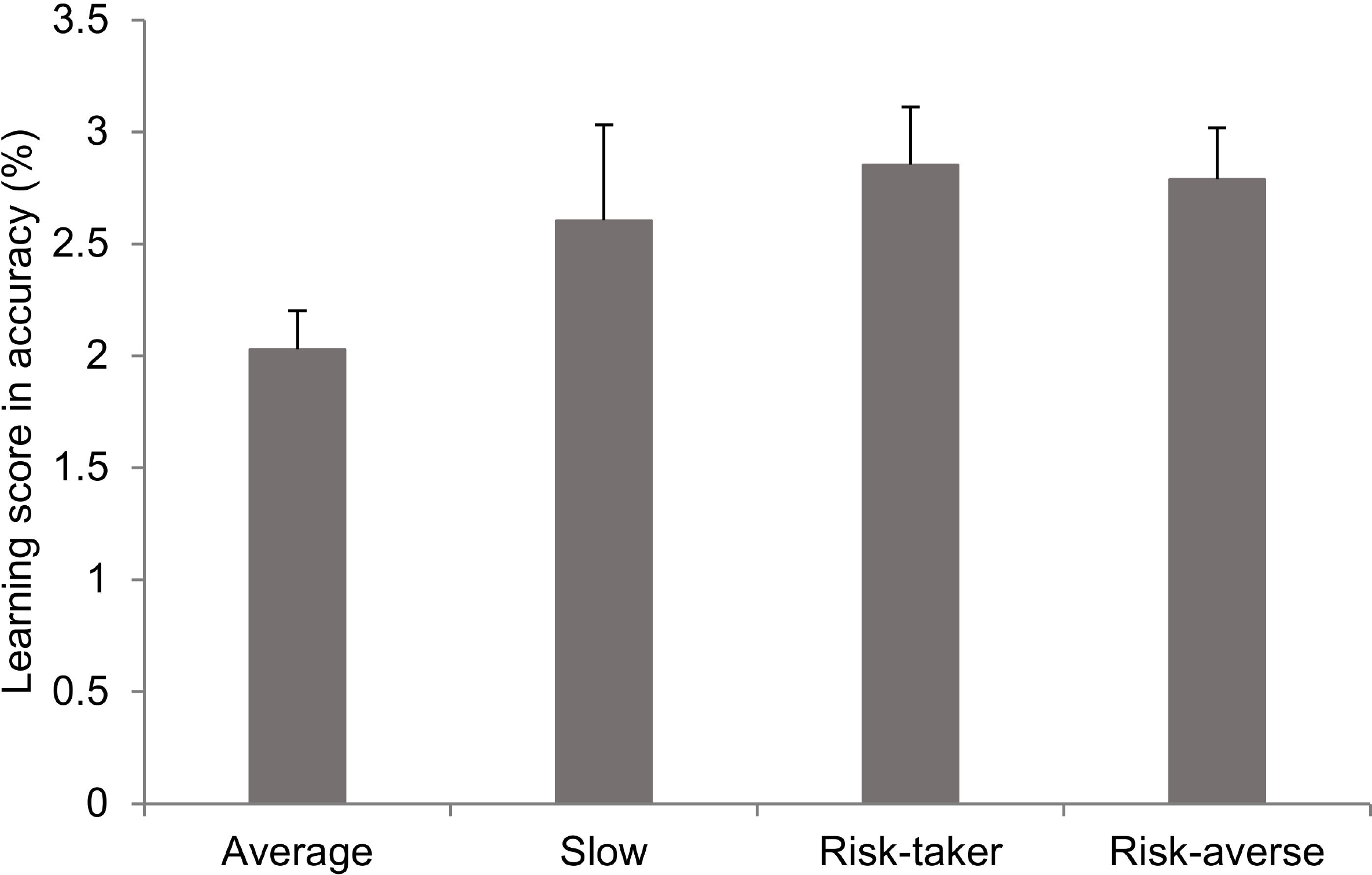
Learning score measure in accuracy for each strategic cluster. Learning score: difference between high-and low-frequency triplets. Error bars denote standard error of mean.

The same ANOVA was performed on *RTs*. In regard to the task-related effects, the entire sample showed probabilistic sequence learning (significant main effects of TRIPLET, *F* (1, 175) = 439.93, *p* < .001, 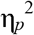 = .715, and EPOCH, *F* (8, 1400) = 268.06, ε = .416, *p* < .001, 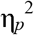 = .605). In addition, participants were increasingly faster on high-than on low-frequency triplets (significant interaction of TRIPLET*EPOCH, *F* (8, 1400) = 30.22, ε = .896, *p* < .001, 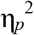 = .147).

Considering the effect of cluster assignment on ASRT RT measures, the main effect of CLUSTER was not significant, *F* (3, 175) = 0.753, *p* = .522, 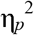 = .013, indicating that overall RT was similar across BART clusters. The non-significant TRIPLET*CLUSTER interaction, *F* (3, 175) = 1.34, *p* = .263, 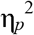 = .022, suggested that probabilistic sequence learning measured by RT did not differ between BART clusters. Similarly, the time course of learning was comparable across BART clusters (non-significant interactions of EPOCH*CLUSTER, *F* (24, 1400) = 0.882, ε = .416,*p* = .550, 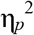 = .015, and TRIPLET*EPOCH*CLUSTER, *F* (24, 1400) = 0.848, ε = .896, *p* = .663,
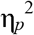 = .014).

In sum, there was evidence for probabilistic sequence learning on both accuracy and RT learning measures in the entire sample. In the case of the accuracy learning measure, this was modulated by participants’ assignment to BART clusters.

## 4. Discussion

This study tested whether implicit probabilistic sequence learning and risky decision making share common variance. To this end, we investigated whether subgroups of participants performing a sequential risk-taking task with probabilistic underlying structure would have been characterized by different sensitivity to statistical regularities measured by an independent probabilistic sequence learning task. According to the results, we successfully identified four different clusters on the basis of usual behavioral measures of the BART. We classified participants as average risk-taking, slowly responding, risk-taker, or risk-averse, respectively. Probabilistic sequence learning was measured by the ASRT task, in which the entire sample, irrespective of cluster assignment, showed significant learning (cf. J. H. Howard, Jr. & Howard, 1997; Nemeth et al., 2010; Nemeth, Janacsek, Polner, et al., 2013). More importantly, we found evidence for greater sensitivity to statistical regularities on the ASRT task in terms of accuracy in the risk-taker and risk-averse subgroups than in participants with average risk-taking for the first time.

We could not have detected association between risky decision making and probabilistic sequence learning if only correlational analysis had been conducted between the individual component measures of the BART and learning scores of the ASRT task. Although our first interpretation of correlational results could have been that no relation was discovered between the two constructs, we have chosen to follow a more detailed description of task-solving profiles with the assumption that this approach might have helped to indirectly reveal the presence of statistical learning in the BART. Since the type of sequence is different in the two tasks in many aspects (predicting the appearance of a stimulus at a certain spatial position vs. predicting the probability of a specific outcome), it has also been possible to assume no relation between the two performance (cf. Goschke & Bolte, 2012). Eventually, our results obtained by clustering suggest that real-world sequential decision making and probabilistic sequence learning are related, at least in some degree. However, as this association could not be directly demonstrated by correlational analysis, we emphasize that further studies should be conducted to support this conclusion with the use of simple or more complex statistical methods. In addition, these studies might directly change the underlying statistical regularities during particular phases of the BART and track whether this manipulation yields a change in performance.

According to the behavioral measures of risk-related performance, participants in the risk-taker group should have been more prone to test the structure of the task as they showed a higher number of risky decisions (see Table 1). Since these participants also showed greater learning in the ASRT task, they might have been inherently, i.e., in a trait-like manner, sensitive to statistical regularities found in both tasks. In addition, they were also found to be less impulsive from a certain aspect as their score was significantly lower on the UPPS Lack of Premeditation scale than that of the other groups (see Table 1, cf. Kaufman et al., 2010). Findings of previous studies suggested that optimal risk taking in the BART was associated with enhanced cognitive capacities shown by neuropsychological and self-report measures as well as by the change in neural activity of the prefrontal cortical areas and along the fronto-striatal network (Bogg, Fukunaga, Finn, & Brown, 2012; Lee et al., 2009). Similarly, it is therefore conceivable that risk-taker participants could have been able to generalize their advantageous cognitive capacities over different but related domains of learning and adaptation; however, this assumption should be further tested and the present results should be replicated in an independent sample.

In the case of risk-averse participants, who also showed greater sensitivity to statistical regularities in the ASRT task, the BART performance essentially differed from that of the risk-taker or average participants. Risk aversion could be considered as default human tendency in uncertain decision making tasks such as the BART (Heilbronner, Hayden, & Platt, 2010; Lauriola et al., 2014), which should be inhibited in order to achieve an optimal or close-to-optimal performance. This notion has further been supported by the decreased risk-taking propensity in previous BART studies testing healthy participants across different versions of the task (Benjamin & Robbins, 2007; Helfinstein et al., 2014; Lauriola et al., 2014; Lejuez et al., 2003; Lejuez et al., 2002; Schonberg et al., 2012; Seaman, Stillman, Howard, & Howard, 2015). According to the results on the change in behavioral performance during the BART, risk-averse participants could have also acquired statistical contingencies during sequential risk taking, but they might have been influenced by other factors such as individual risk preferences, their current emotional or motivational states, experience with the previous balloon or with similar gambling situations, and general problem-solving strategies (Brand et al., 2006; Kardos et al., 2016; Kóbor et al., 2015; Schonberg et al., 2011; Sonuga-Barke, Cortese, Fairchild, & Stringaris, 2015). The latter components also play important role in decision making, and it is unknown to what extent the BART performance reflects differences in probabilistic sequence learning or in these components. A further study manipulating the underlying statistical regularities of the BART could clearly disentangle trait-like sensitivity to statistical regularities (observed in the case of risk-taker participants) and those adaptation mechanisms that enable close-to-optimal behavior on the task according to the experienced response-outcome contingencies. However, it would remain an issue whether probabilistic sequence learning is modulated by the above-mentioned factors at different phases of the task.

Beyond risk-taker and risk-averse participants, the applied clustering method provided the possibility to identify a relatively special subgroup, the slow responders. Slower response time could be related to explorative, more deliberative risk assessment behind decision making processes (Pleskac & Wershbale, 2014), which, in regard to the achieved total score on the BART, might not be the most effective task-solving strategy. This observed pattern could also mirror some aspects of a model-based strategy used by the participants (for the two-system reinforcement learning architecture, see Daw, Gershman, Seymour, Dayan, & Dolan, 2011); however, the latter explanation should be further tested and until then treated with caution as the slowly responding group was the most heterogeneous and the smallest in sample size.

In this study, group differences emerged in the accuracy learning measure but not in the RT learning measure. It has been suggested that accuracy and RT reflect different aspects of probabilistic sequence learning in the ASRT task (S. Song, J. H. Howard, & D. V. Howard, 2007a; Song et al., 2007b). The widening gap between RTs for high-and low-frequency triplets represents mastering the structure of the task via the automatization of predictable responses (J. H. Howard, Jr. & Howard, 1997; Nemeth, Janacsek, & Fiser, 2013). Although RT has been used as the conventional learning measure in the ASRT task, probabilistic sequence learning consists of multiple processes (Nemeth, Janacsek, Király, et al., 2013), and other measures could go beyond the automatization of responses. Namely, accuracy has been a particularly good indicator of prediction errors (Song et al., 2007a), which have also been thought to reflect learning of statistical regularities (J. H. Howard, Jr. & Howard, 1997; Song et al., 2007a; Song et al., 2007b). In particular, as participants gain experience about the underlying statistical regularity of the task (i.e., they learn the high-frequency triplets), they implicitly generate predictions about the likely spatial position of the next stimulus. If the next stimulus is a low-frequency triplet, occurrence of a prediction error is more likely because they expect the high-frequency triplets to a greater extent. As a consequence, overall accuracy also decreases, which pattern has often been reported in probabilistic learning tasks (Curran, 1997; Feeney, Howard, & Howard, 2002; D. V. Howard & Howard, 2001; J. H. Howard, Jr. & Howard, 1997; Schvaneveldt & Gomez, 1998). Thus, in the present study, participants with distinctive risk-taking profiles differed in the prediction-related processes of statistical learning.

This particular finding could originate from the fact that the BART is not a speeded RT task, and in this version, participants had unlimited time to initiate a pumping response or to collect the accumulated reward. Response time variability in the BART has been indicative of different task-solving strategies involving slower, more deliberative decisions and faster, more automatized decisions (Hassall et al., 2013; Pleskac & Wershbale, 2014). Therefore, overall response time, which was merely one of the BART component measures considered in our analysis, might have only been partially related to the RT learning score of the ASRT task. However, no indication was found for this relation here. Nevertheless, to more precisely measure the underlying processes of probabilistic sequence learning, further studies should test both learning measures (accuracy and RT) when investigating the relation of statistical learning and risk-taking behavior.

Another crucial implication of the obtained cluster solution is that a single or a couple of behavioral measures of BART performance might not reliably predict maladaptive risk-taking behavior or the chance to further develop certain psychiatric conditions (e.g., substance abuse/dependence, bipolar disorder, ADHD, etc.). Our results demonstrate that high MAP score does not necessarily indicate excessive risk taking or increased level of impulsivity. Indeed, risk-taker participants collected the largest amount of reward in the BART and outperformed others in a probabilistic sequence learning task, during which they were completely unaware of the acquired regularities. To proceed with these findings, the association between performance on the BART and on the ASRT task should be examined in a concurrent study with clinical populations having atypical fronto-striatal functioning, as their relation has not been clarified in the case of impaired performance.

Although we found evidence for connections between probabilistic sequence learning and risky decision making, the *exact stage* of decision making that is related to probabilistic sequence learning remains uncertain. According to the unified neuroeconomic model of decision making proposed by Sonuga-Barke et al. (2015), decision-making processes involve different psychological stages, which are controlled by distributed and interacting neural circuits. As both implicit and explicit processes affect the different stages of decision making, which, due to the structure of the BART, can be tested separately, further studies should clarify the exact nature of the relation we found here using neuroimaging methods.

Taken together, the present study went beyond the quantification of basic behavioral indices related to BART performance towards a complex characterization of task solving, which more clearly reflected individual differences in risky decision making. Results showed common underlying processes in risky decision making and statistical learning. In addition, we highlighted an adaptive aspect of distinctive risk-taking profiles, which could provide testable assumptions for further neuroimaging studies. Finally, our results could contribute to the refinement of complex neurocognitive models of decision making that is an essential factor in both healthy and impaired daily functioning.

## Author Contributions

AK: basic and advanced data analysis and interpretation, drafting of the work, final approval of the version to be published, agreement to be accountable for all aspects of the work; ÁT: basic data analysis and interpretation, drafting of the work, final approval of the version to be published, agreement to be accountable for all aspects of the work; KJ: design of research, supervision of data acquisition, basic data analysis and interpretation, drafting of the work, final approval of the version to be published, agreement to be accountable for all aspects of the work; ZK: interpretation of results, drafting of the work, final approval of the version to be published, agreement to be accountable for all aspects of the work; CsV: interpretation of results, drafting of the work, final approval of the version to be published, agreement to be accountable for all aspects of the work; DN: supervision of the project, design of research, supervision of data acquisition, interpretation of results, drafting of the work, final approval of the version to be published, agreement to be accountable for all aspects of the work.

